# Functionally Adaptive Structural Basis Sets of the Brain: A Dynamic Fusion Approach

**DOI:** 10.1101/2024.06.18.599621

**Authors:** Marlena Duda, Jiayu Chen, Aysenil Belger, Judith Ford, Daniel Mathalon, Adrian Preda, Jessica Turner, Theo Van Erp, Godfrey Pearlson, Vince D. Calhoun

**Affiliations:** Tri-Institutional Center for Translational Research in Neuroimaging and Data Science (TReNDS), Georgia State University, Georgia Institute of Technology, and Emory University, Atlanta, GA 30303, USA; Department of Psychiatry, University of North Carolina, Chapel Hill, NC, USA; Mental Health Service, San Francisco Veterans Affairs Healthcare System, San Francisco, CA, 94121, USA; Department of Psychiatry and Weill Institute for Neurosciences, University of California San Francisco, San Francisco, CA, 94143, USA; Department of Psychiatry and Human Behavior, University of California Irvine, Irvine, CA, 92697, USA; Department of Psychiatry and Behavioral Health, Wexner Medical Center, The Ohio State University, Columbus, Ohio, 43210, United States; Clinical Translational Neuroscience Laboratory, Department of Psychiatry and Human Behavior, University of California Irvine, Irvine, CA, 92697, USA; Center for the Neurobiology of Learning and Memory, University of California Irvine, Irvine, CA, 92697, USA; Departments of Psychiatry and Neuroscience, Yale University School of Medicine, New Haven, CT, USA; Departments of Psychology and Neuroscience, Georgia State University, Atlanta, GA, 30302, United States

## Abstract

The precise relationship between brain structure and function has been investigated through a multitude of lenses, but one detail that is held constant across most neuroimaging studies in this space is the identification of a singular structural basis set of the brain, upon which functional activation signals can be reconstructed to examine the linkage between structure and function. Such basis sets can be considered “functionally independent”, as they are derived through structural data alone and have no explicit association to functional data. Recent work in multimodal fusion has facilitated a more integrated view of structure-function linkages by enabling the equal contribution of both modalities to the joint decomposition, resulting in components that are independent within modality but co-vary closely across modalities. These existing symmetric fusion approaches thus identify structural bases given an associated functional context. In this work we consider an additional layer of precision to the investigation of structure-function coupling by studying these context-dependent linkages in a time-resolved manner. In other words, we ask which features of brain structure become (or remain) salient given the dynamically changing functional contexts (i.e., dynamic functional connectivity states, task structure, etc.) the brain may pass through during a given fMRI scan. We introduce “dynamic fusion”, an ICA-based symmetric fusion approach that enables flexible, time-resolved linkages between brain structure and dynamic brain function. We show evidence that temporally resolved, and functionally contextualized, structural basis sets can accurately reflect dynamic functional processes and capture diagnostically relevant structure-functional coupling while detecting nuanced functionally-driven structural components that cannot be captured with traditionally computed structural bases. Lastly, differential analysis of component stability across repeated scans from a control cohort reveals organization of static and dynamic structure/function coupling falls along unimodal/transmodal hierarchical lines.

## Introduction

Over the last 50 years, advances in neuroimaging have produced a host of techniques capable of capturing detailed signatures of brain structure and function. Along with its potential for precision diagnostics in psychiatric and neurological conditions, the rise of neuroimaging has been instrumental in furthering investigation into one of the most fundamental questions in neuroscience: what is the relationship between brain structure and function? To this end, a class of approaches have been developed under the umbrella known as multimodal fusion, which aim to optimally integrate complementary data across various neuroimaging modalities. Multimodal fusion spans a wide range of techniques capable of revealing hidden linkages between imaging data types, and these techniques can be broadly divided into two categories: asymmetric fusion approaches, wherein data from one modality are used to constrain or influence the analysis of another modality, and symmetric fusion approaches, wherein each imaging modality contributes equally to the combined analysis, fully leveraging the joint information across datasets (V. D. Calhoun & Sui, 2016; Sui et al., 2012).

In utilizing symmetric multimodal fusion to study the coupling between brain structure and function, there arises the common challenge of dealing with dimensional incongruence between “static” measures of brain structure and functional neuroimaging datatypes that include an additional temporal dimension. Historically, this has been addressed by heavily summarizing over the temporal dimension of the functional time series by computing measures such as static functional network connectivity (sFNC) (DeRamus et al., 2022) or amplitude of low-frequency fluctuation (ALFF) (Lottman et al., 2018), which does achieve dimensional compatibility of the data types for fusion, but also effectively removes the rich temporal information of the functional data. Some data driven approaches including parallel ICA (Vergara et al., 2014) group ICA + ICA (Qi et al., 2022) have been developed to handle mismatched data dimensionality, however these have not yet been expanded to fully leverage functional dynamics. With the surge in focus on functional dynamics over the last decade, the field has begun to recognize the significance of short-scale variation in connectivity (particularly during rest) and has pushed to preserve more temporal information in the analysis of fMRI data. A common output of dynamic functional network connectivity (dFNC) analysis is the identification of a small set of dFNC states or patterns that subjects may transition within and between during rest, often derived via k-means clustering (Allen et al., 2014). Several multimodal fusion studies have operated on the level of dFNC states by concatenating subject-level average connectomes across all states into a single FNC “mega-vector” in the fusion design (Abrol et al., 2017; Duda et al., 2022), thus enabling the inclusion of dynamic information into the multimodal fusion without increasing dimensionality of the functional inputs. Symmetric fusion approaches, such as multiset canonical correlation analysis with joint ICA (mCCA + jICA), were then applied to find hidden linkages between the concatenated dFNC and structural data, and functional patterns specific to each dFNC state were reconstructed post-hoc. While this approach enables some interpretation of state-specific linkages to structure, the resultant couplings are rather rigid as the concatenated dFNC components necessarily share a single loading parameter linking to a single structural component, imposing the (likely too strict) assumption that linkage to structure is identical across all dFNC states. Other joint approaches have investigated functionally informed structural basis sets (Plis et al., 2018) as well as individual linkages between specific intrinsic functional networks and structure (Khalilullah et al., 2023), but thus far there has been no framework for systematically identifying flexible structural basis sets based on linkages to time varying brain function.

On the flip side, widely used asymmetric fusion approaches are not bound by the same dimensionality constraints as symmetric fusions, at the cost of being less informative about the joint relationship between modalities. One particular line of research that has recently emerged and can be classified under the umbrella of asymmetric fusion is the study of “structural brain modes”. In short, structural graphs (which can be derived as white matter structural connectivity matrices via tractography analysis on diffusion MRI (Griffa et al., 2022) or maps of cortical surface geometry (Pang et al., 2023)) are subjected to eigenmode decomposition to obtain a singular structural basis set representing the so-called “structural harmonics” of the brain. These structural modes can be linked to brain function by projecting the fMRI signal at each timepoint as weights onto the discovered structural manifold. Such an approach does enable some flexibility in the linkage of structure to function over time through modification of the weighted mixture of the structural modes, however the rigid definition of only one structural basis set derived without explicit linkage to functional data assumes that the full realm of complex brain functions can be sufficiently described by a singular set of structural modes.

It has long been known that functional brain activity and connectivity vary meaningfully over both long (across the lifespan) (Battaglia et al., 2020; Baum et al., 2020) and short (seconds to minutes) (V. Calhoun et al., 2010; V. D. Calhoun et al., 2014; Chang & Glover, 2010; Lurie et al., 2020; Preti et al., 2017) timescales, however the potential for similar temporal variation in structure-function couplings, especially at shorter timescales, has thus far been understudied. Most prior work has focused on either regional (Fukushima et al., 2018; Fukushima & Sporns, 2020) or temporal (Preti & Van De Ville, 2019; Vázquez-Rodríguez et al., 2019) fluctuations in structure-function coupling, but attempting to study simultaneous spatiotemporal fluctuation patterns of structure-function linkages is a relatively novel concept. One such work utilized multilinear regression to link regional measures of instantaneous functional connectivity to their structural connectivity profiles derived from diffusion MRI (dMRI) as an estimate of time- and region-resolved structure-function coupling (Liu et al., 2022). This study found that dynamic coupling patterns reflect cortical hierarchies, showing the strongest spatiotemporal fluctuations in the insula and frontal eye fields and the least in the unimodal and transmodal cortices. This atlas-based approach represents a critical initial dive into the space of temporally-evolving structure-function coupling at the level of data integration, and continued investigation into this space with advanced approaches like multimodal fusion promises to yield even deeper insights in this emerging domain of research by enabling joint estimation of data-driven structure-function linkages at the voxel level.

Here, we introduce an ICA-based symmetric fusion approach that allows for the identification of a temporally adaptive basis set that is inclusive of both structure and function, which we term “dynamic fusion”. We wish to be explicit in the fact that we **do not claim** that neuroanatomical structure is physically changing on short timescales. Rather, we explore the concept that a single basis set derived from structural data alone may not provide the best representation of the highly complex interplay of brain structure required for the full range of human brain function. We advocate for the creation of structural features (i.e. basis sets/latent representations) in a way that is informed by the dynamically changing functional contexts that individuals oscillate through across the duration of a functional scan. Consider, as an intuitive example, the swinging pendulum system. The system has finite set of physical properties that are unchanging during the observation period (akin to the structural properties of the brain that are unchanging during the course of an MRI scan); however, depending on the functional aspect of interest in the model (e.g., oscillatory motion or energy dissipation), different physical (i.e. “structural”) aspects of the system become salient (e.g., angular position, angular velocity and pendulum length for oscillatory motion, and velocity, surface area, coefficient of drag and friction at pivot for energy dissipation), and thus different structural basis sets are defined for the same system. In the same way, our dynamic fusion approach derives multiple structural basis sets for the same physical brain system that are each best suited to describe the different functional “states” considered across time. In other words, what we derive are temporally evolving patterns of structure-function *coupling*, rather than short-scale changes in physical brain structure itself. The symmetric aspect of the mCCA + jICA algorithm in our dynamic fusion framework also enables both modalities to equally contribute to the joint solution, so though the variability in component maps across fusions is driven by functional influence of each dFNC state by virtue of holding the structural inputs constant, the solution for each fusion individually is based on the shared co-variance between structure and that specific functional context. In this way, dynamic fusion can uncover more nuanced, temporally evolving structure-function relationships than traditional methods.

The following paper highlights three key results. First, inter-state cross-fusion comparisons revealed that structural components exist on a spectrum from high to low cross-fusion stability, which can be interpreted as less functionally influenced (i.e., “static”) to more functionally influenced (i.e., “dynamic”). Second, we show dynamic structural components exhibit the strongest group differences in component loading parameters in a schizophrenia (SZ) vs. control cohort, suggesting these variations in component maps are not arbitrary, but are driven by meaningful linkages to function. Third, we show that the organization of the brain into regions of highly static vs. highly dynamic structure-function coupling largely falls along unimodal/transmodal hierarchical lines. These results indicate that dynamic fusion enables flexible linkages between structure and time-evolving function which are related to changing neural contexts, and suggest that the functionally influenced dynamic components may capture clinically relevant structure-function linkages that would be otherwise missed by standard approaches. In addition to fMRI/sMRI fusion experiments, we show similar results with dMRI as structural inputs to the dynamic fusion pipeline indicating measures of both white and gray matter show evidence for time-varying coupling to function.

## Methods

### Data & Preprocessing

We analyzed functional, structural, and diffusion MRI from the HCP 1200 (Van Essen et al., 2013) (N = 833 subjects, average age = 28.7 years) and FBIRN (Keator et al., 2016) (N = 310 subjects, N_SZ_ = 150 subjects [114 male, average age = 38.8 years], N_CON_ = 160 subjects [115 male, average age = 37.0 years]) datasets. All data were collected with informed consent and approved by the Institutional Review Boards at the participating institutions. Processing pipelines are detailed below:

#### fMRI

In the FBIRN dataset, resting state fMRI data were preprocessed utilizing the statistical parametric mapping (SPM12, http://www.fil.ion.acl.ac.uk/spm/) toolbox within MATLAB 2019. Initially, the first five scans were discarded to allow for signal equilibrium and to allow for participants to adapt to the scanner noise. We then applied rigid body motion correction using the mean scan/frame as reference to address any head motion by the subjects, followed by slice timing correction using the median slice as reference to account for timing discrepancies in slice acquisition. The fMRI data were subsequently transformed into the standard Montreal Neurological Institute (MNI) space using an echo-planar imaging (EPI) template, with slight resampling to 3 x 3 x 3 mm^3^ isotropic voxels. The resampled fMRI images were then smoothed with a Gaussian kernel, achieving a fill width at half maximum (FWHM) of 6 mm. For the HCP 1200 release data, we downloaded the preprocessed data from online, normalized them to MNI space and resliced them to the same spatial resolution (3 x 3 x 3 mm^3^) using SPM12. More detailed in terms of preprocessing on HCP data can be found online (http://www.humanconnectomeproject.org/data/). Subjects with head motion larger than 3° rotations or 3 mm translations or with mean framewise displacement (FD) larger than 0.3 mm were excluded.

Spatially constrained ICA was applied using the GIFT toolbox (https://trendscenter.org/software/gift/) (Iraji et al., 2021) using the NeuroMark_fMRI_1.0 template (Du et al., 2020) to extract subject-level spatial maps and for each of the 53 intrinsic connectivity networks (ICNs) defined in the template, as well as their respective activation time courses. Dynamic FNC (dFNC) analysis of fMRI differed between the HCP and FBIRN datasets in an effort to explore consistency of the results of our dynamic fusion approach relative to upstream pipeline design choices.

In FBIRN, time and frequency resolved FNC patterns were computed from ICN time courses using the filter-bank connectivity (FBC) approach (Faghiri et al., 2021). Briefly, FBC utilizes a filter bank, i.e., an array of systems used to filter a time series into different frequency bands (typically non-overlapping and spanning the full frequency spectrum of the data), which enables estimation of FNC within a given frequency range. We designed our filter bank to contain 10 Chebyshev type-2 infinite impulse response filters that evenly cover the full frequency spectrum of the fMRI time series (0.00 – 0.25 Hz). K-means clustering identified six distinct states with unique connectivity signatures and spectral occupancy across frequency bands. For more detail on the FBC approach and state clustering used here see (Duda et al., 2022).

In HCP, we utilized the REST1_LR and REST2_LR fMRI time series to perform repeated experiments within the same subject set and scan type. We computed dFNC in resting state fMRI from ICN time courses using a sliding window Pearson correlation (SWPC) approach outlined in (Damaraju et al., 2014), with the exception that the window size used in our analysis was 10 TR (∼15s; TR = 0.72s). We chose a 10 TR window to keep consistency between the window size in HCP and FBIRN analyses. K-means clustering in resting state data revealed five dFNC states, and subject-average connectomes were computed for each state for dynamic fusion.

#### sMRI

T1-weighted sMRI images were preprocessed using the DARTEL tool from SPM12 within the MATLAB 2019 environment. Image registration, bias correction, and tissue clarification were performed using the unified model. DARTEL was then employed to estomate the gray matter volume (GMV) images, which were resliced to a resolution of 1.5 mm and smoothed with a Gaussian kernel with FWHM = 10 mm.

#### dMRI

The processing of dMRI data in HCP involves a series of steps using FSL (www.fmrib.ox.ac.uk/fsl), ANTs (Avants et al., 2009), and custom-built software. First, the datasets were collected with phase reversed blips, resulting in images with distortion in opposite directions. To address it, volumes obtained at b0 (b=5 s/mm^2^, with phase encoding in the left-right (LR) and right-left (RL) direction) were extracted from the entire set of 576 volumes for each dataset (comprising 3 LR runs with 95, 96 and 97 directions respectively, and 3 RL runs with corresponding directions). These b0 volumes were preselected by identifying the least-distorted pair, which was then used with topup (FSL 6.0)(Andersson et al., 2003) to estimate the susceptibility-induced off resonance field. The estimated fieldmap was used to unwarp the phased-encoded dMRI images. Next, the dMRI volumes underwent correction for eddy current distortions and head movement using eddy_gpu (FSL 6.0), which incorporated advanced motion-induced signal dropout detection and replacement (Andersson & Sotiropoulos, 2016).

Subsequently, Fractional Anisotropy (FA) maps were estimated from the diffusion tensor model using dtifit (FSL 6.0). The FA maps then underwent automated noise inspection, where any detected noise spots were padded with interpolation, and all FA values were adjusted to fall within the range of 0 to 1.

Following this, the FA maps were registered to the Montreal Neurological Institute (MNI) standard space through nonlinear registration performed by ANTs. Any images exhibiting excessive motion, signal dropout, or excessive noise were excluded from subsequent analysis (Geenjaar et al., 2023; Wu et al., 2015).

### Dynamic Fusion

We used multi-set canonical correlation analysis followed by joint independent component analysis (mCCA + jICA) to perform fusion of both static (GMV/FA maps) and temporally resolved (SWPC/FBC states) neuroimaging features. The combined mCCA + jICA model is designed to allow for the identification of both strongly and weakly correlated joint components that are also independent from one another. This is achieved by employing mCCA in the first step to generate flexible linkages between the modalities, and subsequently applying jICA to ensure independence between components. The ability to identify flexible (i.e., both strong and weaker) linkages between modalities is essential for the application to dynamic fusion, in which we are searching for nuanced differences in the joint information between structure and dFNC as a function of time, thus this requirement informed our design choice of employing mCCA + jICA as the symmetric fusion algorithm. The mCCA + jICA framework is defined under the assumption that a multimodal dataset, *X_k_*, is a linear mixture of *m* sources (S_k_) mixed by non-singular matrices (*A_k_*), here, *k* = (1,2) and model order *m* = (10, 15) sources/components. The effective mCCA + jICA framework can be defined as *X_k_* = (*D_k_W^-1^)S_k_*, where modality-specific mixing matrices are defined as *A_k_* = *D_k_W^-1^*. Further details can be found in (Sui et al., 2011, 2013).

The key novelty in dynamic fusion is the concept of performing multiple data fusions which are linked together. In short, dynamic fusion generates functionally adaptive basis sets of brain structure by conducting multiple fusion experiments linking a static neuroanatomical measure (e.g., GMV map) to various distinct functional measures that represent temporally evolving functional patterns (Fig 1). This can in theory be applied at various levels of functional granularity, but since resting state time courses are not temporally aligned across subjects, window-level dynamic fusion would not be appropriate as the underlying functional contexts/cognitive processes would not be fixed across subjects within each fusion. Thus, to enforce some level of temporal and functional alignment in the resting state fMRI experiments, we perform state-level dynamic fusion wherein structural measures (either GMV or FA maps) are separately fused with subject-level state-average FNC matrices. All dynamic fusion experiments resulted in a set of structural (GMV or FA) components (model order = 15 components) optimized to each time-resolved FNC pattern independently. Cross-fusion (i.e., inter-state) comparisons of these structural components across experiments revealed structural manifolds unique to each distinct functional basis (i.e., GMV/FA components related to frequency-specific functional connectivity dynamics), as well as some that were identified across multiple fusions, indicating a “static” structural component. In addition to the inter-state fusion experiments, we conducted intra-state (i.e., same-state) cross-fusion analyses to serve as baselines. For each subject, timepoints corresponding to each dFNC state were randomly split into two equal sets and subject-level average FNC was computed using the only the timepoints in each set. Each split-half FNC matrix was used as a functional input to separate fusions and cross-fusion correspondence was computed between fusions from the *same* state, thus assessing component stability across fusions where the functional inputs were not identical but derived from the same functional context.

**Fig 1.**
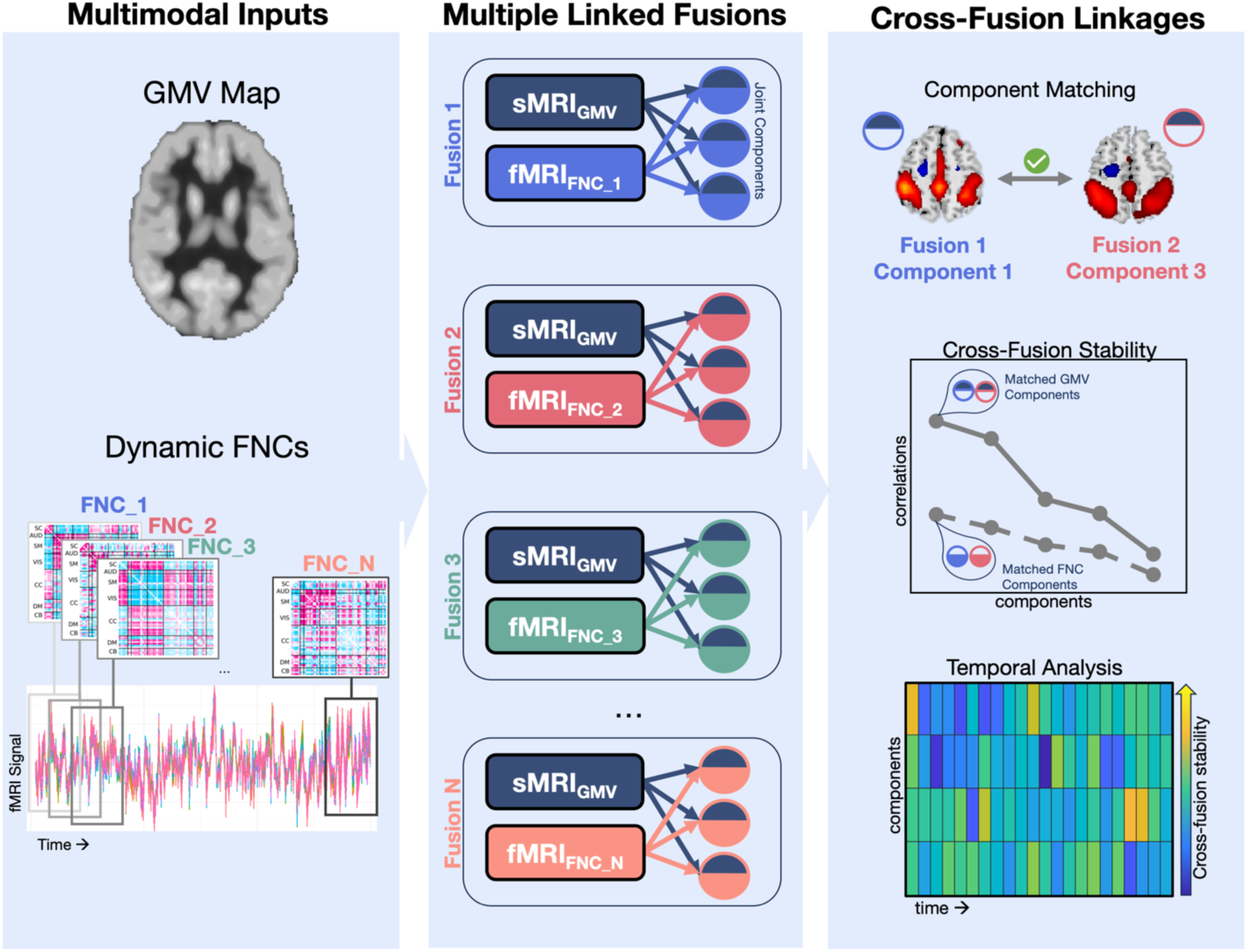
Schematic of dynamic fusion experimental design. Dynamic fusion links a single structural measure (i.e., GMV maps) to dynamic functional measures (i.e., windowed FNC matrices or subject average state connectomes) via multiple linked fusion experiments. Under the dynamic fusion framework, these can be performed as separate fusions to extract multiple functionally influenced structural basis sets. Post-hoc cross-fusion analysis is performed to assess the stability of both the structural and functional components derived across the linked fusions, as well as their temporal evolution over time.

## Results

### Cross-State Fusions Show Significant Functional Specification of Both Functional and Structural Component Maps at Rest

In both the HCP and the FBIRN datasets, we performed inter-state dynamic fusion (as well as intra-state fusion baselines) of GMV maps with different metrics of dFNC for each dataset (five FNC states in HCP and six FBC states in FBIRN). Fig. 2 illustrates the cross-fusion stability of structural (left) and functional (right) component maps for both FBIRN (top) and HCP (bottom) experiments. FBIRN analyses included six intra-state pairs for baselines and 15 inter-state pairs for experimental results, and HCP included ten intra-state pairs (five each for REST1 and REST2 scans) for baselines and 20 inter-state pairs (ten each for REST1 and REST2 scans) for experimental results.

**Fig. 2.**
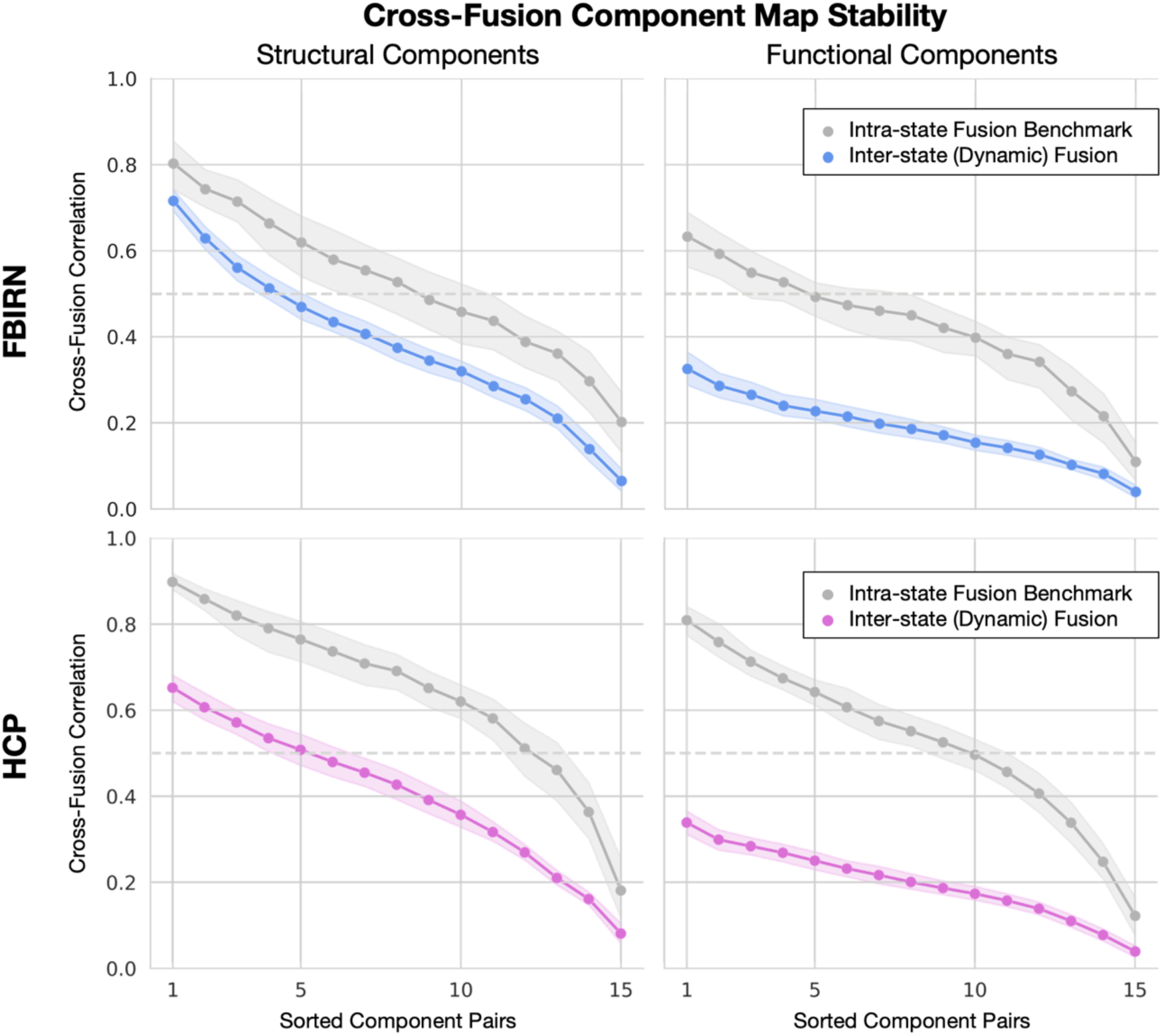
Cross-fusion stability of structural and functional component maps in both FBIRN (top) and HCP (bottom) dynamic fusion experiments across all sorted component pairs (model order = 15). Points and error bands correspond to the means and 95% CI, respectively, of all the cross-fusion comparisons of each type in each dataset (FBIRN: 6 intra-state, 15 inter-state; HCP: 10 intra-state, 20 inter-state). Static/dynamic threshold r = 0.50 shown in dotted line. Note that structural components are generally more stable across fusions (as expected), but interestingly a small subset of structural components are highly stable and another larger subset are more ‘dynamic’ that is they vary across fusions based on the context provided by the linked fMRI data.

Across both datasets, intra-state baselines exhibit high stability across fusions in both structural and functional components, providing the expected cross-fusion correspondence of component maps due to random variation within a constant functional state or context. Experimental inter-state results showed significantly lower cross-fusion correspondence/stability (p < 0.05 – see Supplemental Tables S1 and S2), or in other words, significant functional specialization of component maps in both structural and functional fusion outputs. Given the structural inputs to all cross-fusion comparisons was identical, paired with the significantly lowered cross-fusion correlations compared to baselines, we can conclude that these component map variations, specifically structural components, are driven by meaningful changes in functional patterns and thus meaningful variations in state-specific structure-function coupling.

Overall, we observed relatively low inter-state correspondence for the functional components which was expected as each of the functional inputs (dFNC states) have unique connectivity signatures, and, in the case of the FBIRN experiments, are even found in distinct frequency bands. Though the GMV components exhibited higher stability overall, we observed varying degrees of correspondence between best-matched GMV components across independent fusion experiments, even though the structural inputs were identical across each fusion. Specifically, we found fairly high cross-fusion correspondence (|r| > 0.5, shown in dotted lines in Fig. 2) in only the first few sorted component pairs, followed by a fairly steep drop-off of component correspondence, indicating little overlap between a majority of the resultant components. These consistent results across the two datasets suggest most of the structural components identified via mCCA + jICA are significantly functionally influenced (i.e., “dynamic”), while a small subset are less functionally influenced (i.e., “static”). This result further demonstrates the ability of symmetric fusion approaches like mCCA + jICA to identify truly joint linkages in the data, as the changes in the functional inputs resulted in considerably different structural outputs, despite accounting for a much smaller portion of the entire dimensionality of the original input data. Moreover, our findings underscore the importance of enabling flexible linkages between structure and changing function, as arbitrarily imposing a limited structural basis would obfuscate the differential structure-function linkages that we have identified via dynamic fusion.

### Significant Schizophrenia/Control Group Differences were identified in both Static and Dynamic Components

For each of the 90 components derived across the six linked FBIRN state-level dynamic fusion experiments (model order = 15), we computed group differences between the structural component loading parameters using a 2-sample t-test (p-values FDR corrected for multiple comparisons) and defined each component as “static” or “dynamic” (i.e., functionally uninfluenced or functionally influenced) based on the average cross-fusion correlation > or < |0.5| across all pairs of cross-fusion experiments, respectively (Fig. 3 & Supplemental Table S3). The cutoff r = |0.5| was chosen as the cutoff as it was both the midpoint of the possible range of values (0 – 1.0) and the true range of values (0.0002 – 0.95) that we observed in the data. We found 9/17 static components exhibited significant group loading differences, as well as a majority (40/73) dynamic components. We find the significance levels of many of the top dynamic components was directly on par with those of the static components; in fact, the most significant group difference in structural loadings was found in one of the most dynamic components (average cross-fusion correlation = 0.090). These results further highlight the utility of functionally influenced dynamic structural components in capturing clinically and diagnostically relevant features of structure-function coupling that are otherwise missed in the static components. In Figures 4 and 5, we visualize the joint sources and highlight group effects for some of the most significant components from both the dynamic and static sets.

**Fig. 3.**
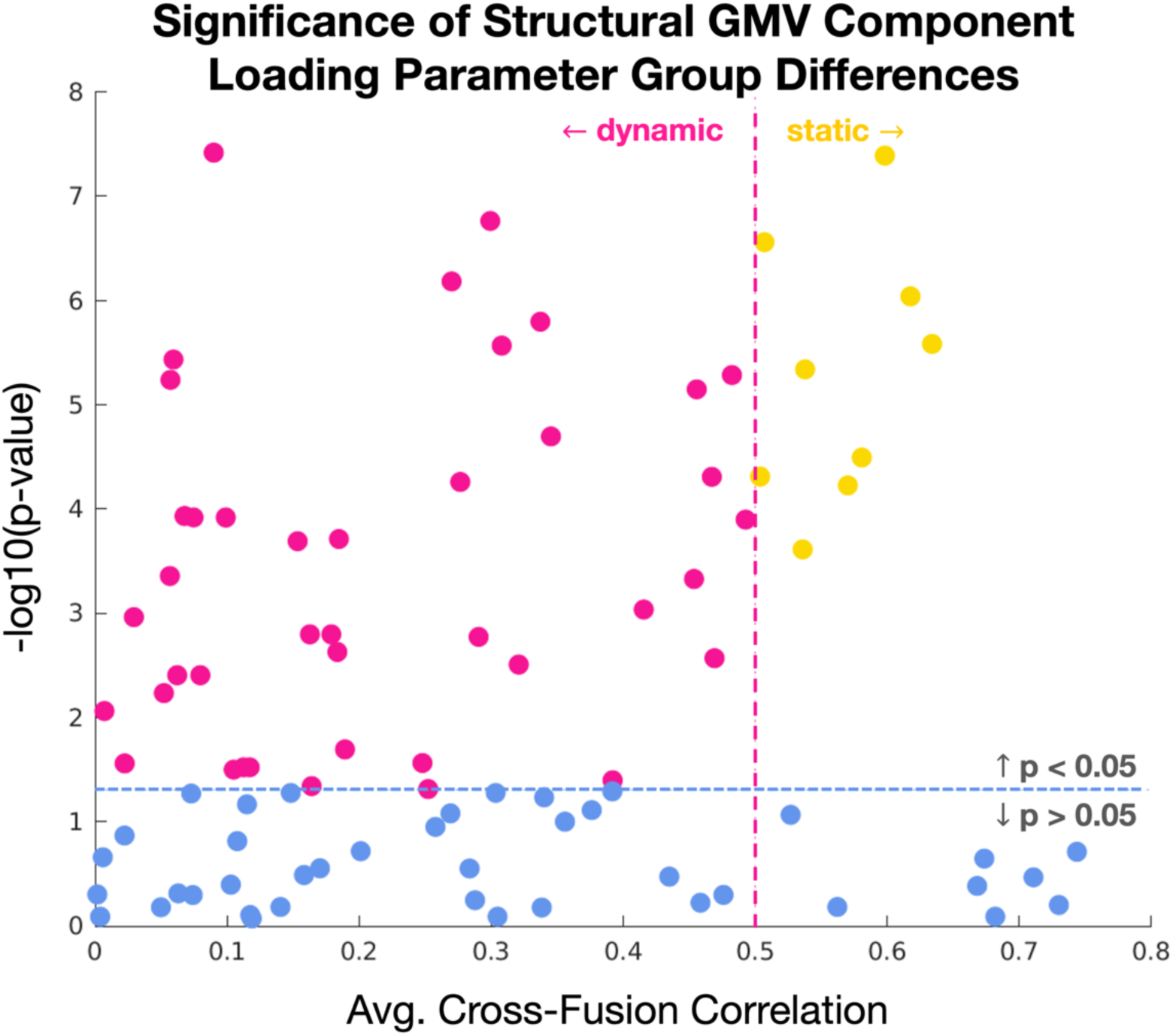
Significance of SZ/control group differences of GMV component loading parameters vs. average cross-fusion correlation of best-matched component maps for each of the 90 components generated across six state-level dynamic fusion experiments. (All p-values FDR corrected for multiple comparisons).

**Fig. 4.**
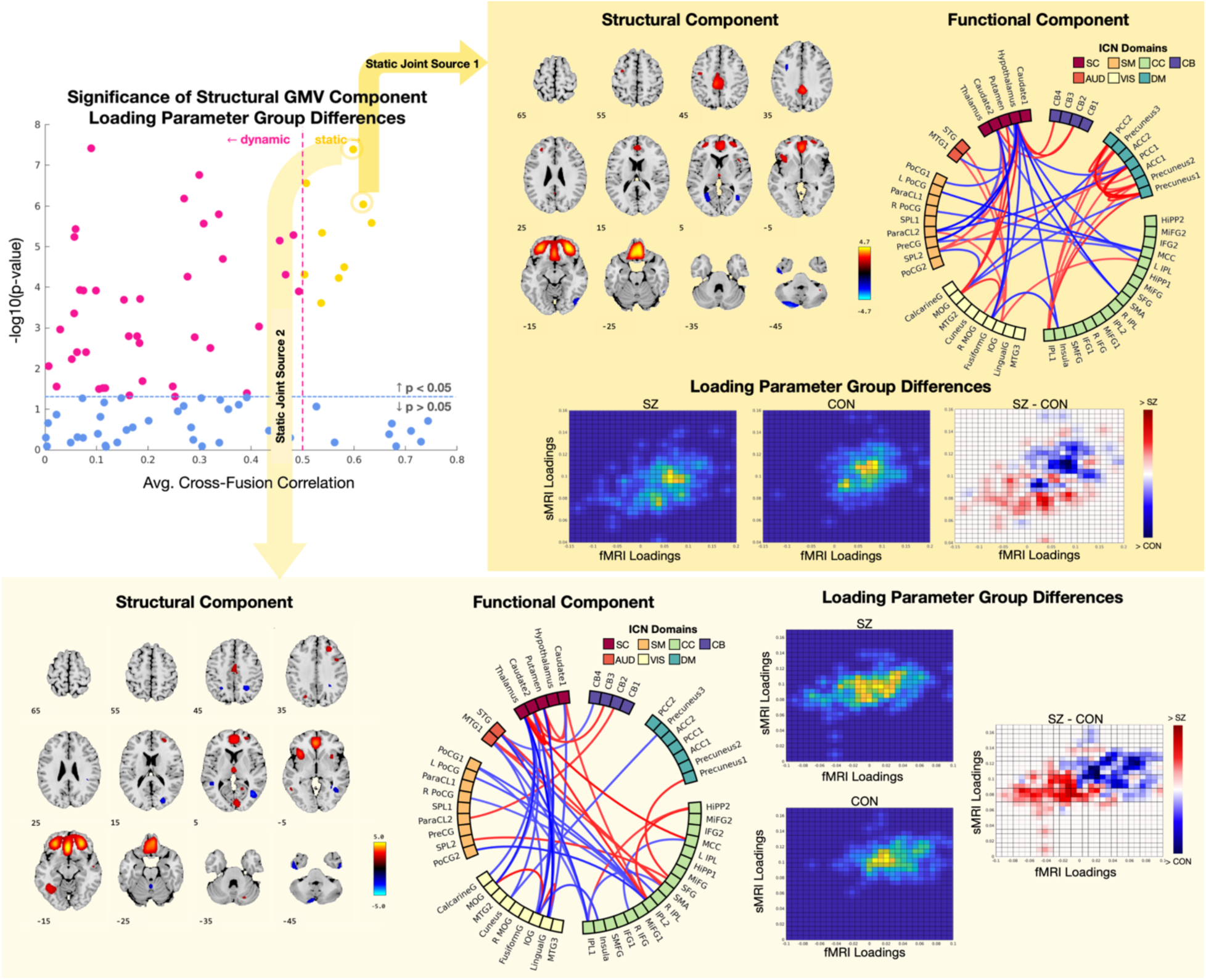
Joint sources of paired static components with significant group differences in loading parameters. Static Joint Source 1 was derived from dynamic fusion to a low-frequency state (State 1), while Static Joint Source 2 was derived from dynamic fusion to a mid-frequency state (State 6). Structural component maps are highly similar, both exhibiting peaks in cingulate and orbitofrontal regions, while linked functional components show little overlap. See (Duda et al., 2022) for more details on FBC states. Joint histograms show where subjects lie in the overall distribution of structural (sMRI) and functional (fMRI) loading parameter space. Abbreviations: subcortical (SC), auditory (AUD), sensorimotor (SM), visual (VIS), cognitive control (CC), default mode (DM), cerebellum (CB), schizophrenia (SZ), control (CON).

**Fig. 5.**
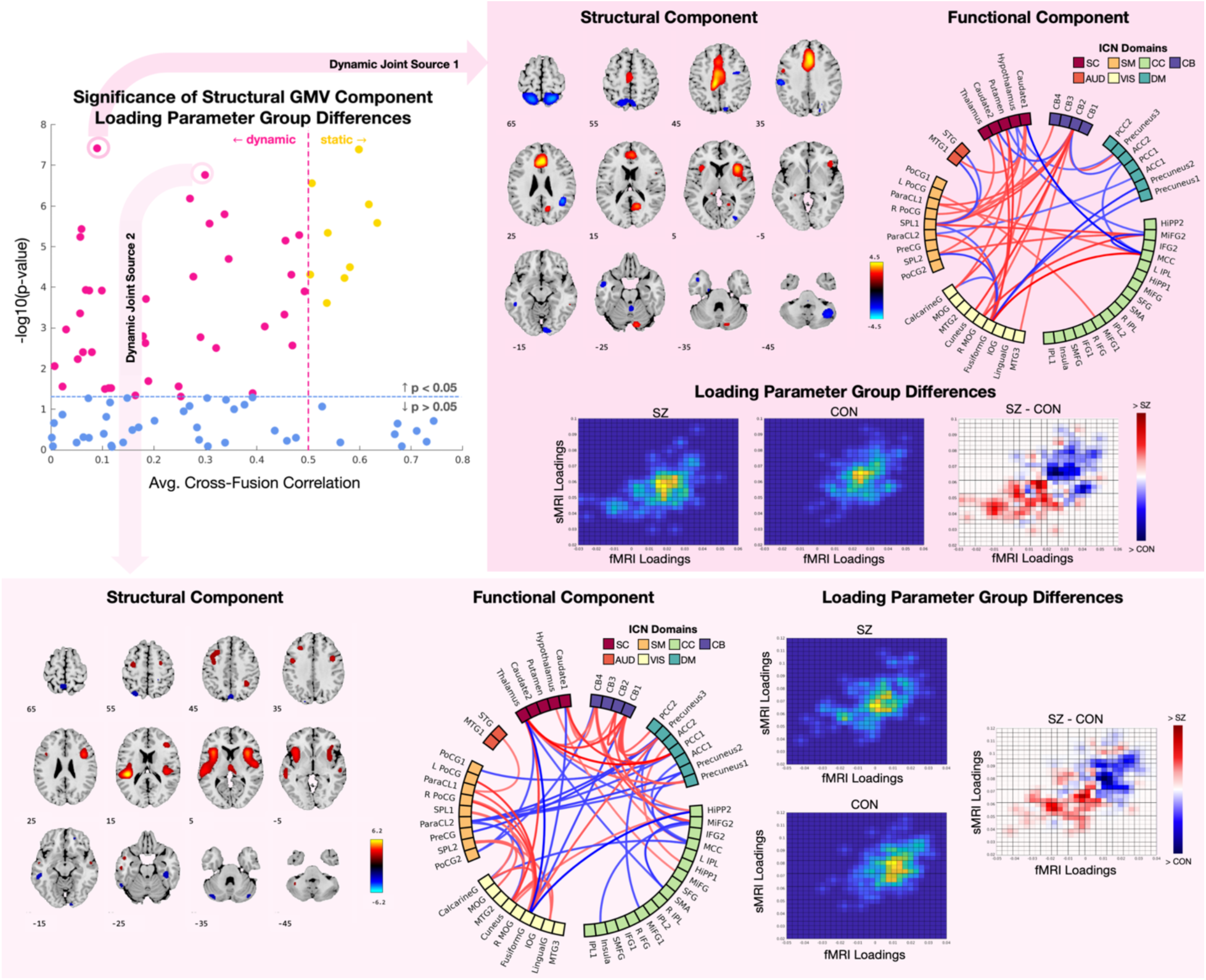
Joint sources of two exemplar dynamic components with significant group differences in loading parameters. Dynamic Joint Source 1 was derived from dynamic fusion to a CON-dominant mid-frequency state (State 4) and Dynamic Joint Source 2 was derived from dynamic fusion to an SZ-dominant high-frequency state (State 6). The structural map for Dynamic Joint Source 1 shows peaks in the cingulate cortex and superior parietal lobule, while the structural map for Dynamic Joint Source 2 shows peaks in inferior parietal cognitive control regions as well as the insula. Joint loading histograms show tight concentration of both SZ and CON subjects, and strong separations on both the structural and functional component axes, indicating strong contributions of both the structural and functional data to the joint sources. See (Duda et al., 2022) for more details on FBC states. Abbreviations: subcortical (SC), auditory (AUD), sensorimotor (SM), visual (VIS), cognitive control (CC), default mode (DM), cerebellum (CB), schizophrenia (SZ), control (CON).

Joint sources for a pair of static components exhibiting significant group differences in loading parameters are shown in Fig. 4. Static Joint Source 1 was derived from fusion with functional data from a low-frequency state (State 1 in (Duda et al., 2022)), and exhibited an average cross-fusion correlation of 0.618 as well as group differences in GMV component loading parameters at a significance level p = 9.14e-07. Static Joint Source 2 was derived from fusion with functional data from a mid-frequency state (State 5 in (Duda et al., 2022)), and exhibited an average cross-fusion correlation of 0.598 as well as group differences in GMV component loading parameters at a significance level p = 4.08e-08. Though both derived from fusions with notably different functional contexts, as underscored by the lack of overlap in most edges in the functional components for each joint source, the structural component maps are highly similar across the fusions (r = 0.738), with peaks in cingulate and orbitofrontal regions. These paired sources illustrate structural components that are not strongly influenced by their link to function, which can be understood as “static” or “global” structural components. This is also reflected in the joint histograms depicting loading parameter group differences – in Static Joint Source 1 the SZ and CON groups are more readily separable across the structural loading axis and more overlapped across the functional loadings axis, suggesting an overall weaker influence of the functional inputs compared to the structural inputs to the entire joint source.

We highlight two dynamic joint sources in Fig. 5. Dynamic Joint Source 1 was derived from a CON- dominant mid-frequency state (State 4 in (Duda et al., 2022)) and Dynamic Joint Source 2 was derived from dynamic fusion to an SZ-dominant high-frequency state (State 6 in (Duda et al., 2022)). Dynamic Joint Source 1 exhibits peaks across the anterior cingulate cortex, insula, and superior parietal regions, has an average cross-fusion correlation of 0.0.90 and group differences in structural component loading params at the significance level p = 3.82-08, making it among one of the most dynamic (i.e., most functionally influenced/functionally specialized) components, as well as the component with the most significant group differences in structural loading parameters. Dynamic Joint Source 2 exhibits peaks in cognitive control regions, including the insula and inferior parietal lobule, has an average cross-fusion correlation of 0.299 and exhibited strong group differences in structural component loading params at the significance level p = 1.73e-07. As expected, the functional components for each dynamic joint source are unique, but some interesting similarities do appear – for example the fusiform gyrus (FFG) within the visual domain as well as CB2 in the cerebellum serve as functional hubs in both functional components. Dynamic Joint Source 1 is marked by positive connections between VIS and CB, SC, and CC regions, as well as CB-SM, and both FFG-DM and SC-CC anticorrelations. Dynamic Joint Source 2 is marked by negative edges connecting SM-DM regions, as well as between FFG and SC, CB and CC regions, and positive correlations between CB and DM, CC and SC regions, particularly caudate and thalamus. Further examination of the dual histograms of loading parameter group differences, we observe tighter concentration of SZ and CON subjects within both the structural and functional loading parameter spaces than was observed for the static sources, as well as more distinct separability of the groups in both the structural and functional parameter axes, suggesting a stronger contribution of both inputs into the joint solution.

### Organization of Static/Dynamic Structure/Function Coupling in Healthy Brains Aligns to Unimodal/Transmodal Hierarchical Segregation of Cortex

After establishing both the cognitive relevance (Fig 2) and clinical utility (Figs 3-5) of joint sources derived via dynamic fusion, we interrogated the organization of strongly static or strongly dynamic regions in the brain. Thus, we applied our dynamic fusion pipeline in both HCP REST1_LR and REST2_LR fMRI time series (henceforth termed Rest1 and Rest2), collected with the same scan parameters on the same subjects on both their initial and return study visits (Van Essen et al., 2013). In both experiments, 5 dFNC states were derived via k-means and model order = 15 were used for fusion, resulting in 75 joint sources each for both Rest1 and Rest2. As was done in the FBIRN analysis, greedy cross-fusion matching and correlation of structural component maps was performed separately in both experiments, and components were ranked and labeled as “dynamic” or “static” according to their stability score (i.e., average cross-fusion correlations) < or > 0.5, respectively. After filtering of spurious components, we selected the top 10 most static (i.e., highest stability scores) and top 10 most dynamic (i.e., lowest stability scores) components. For each set of static/dynamic components, component maps are aggregated as follows: first, component maps are thresholded for significance (|z| >= 2), then |z| maps are normalized to [0, 1], weighted by stability score (or 1-stability score in the case of dynamic components), and aggregated by taking the max value at each voxel. For both experiments, we subtract the dynamic map from the static map to obtain composite differential stability maps across the whole brain, and finally average the Rest1 and Rest2 maps to obtain a global view of brain organization into statically/dynamically linked regions (Fig. 6).

**Fig. 6.**
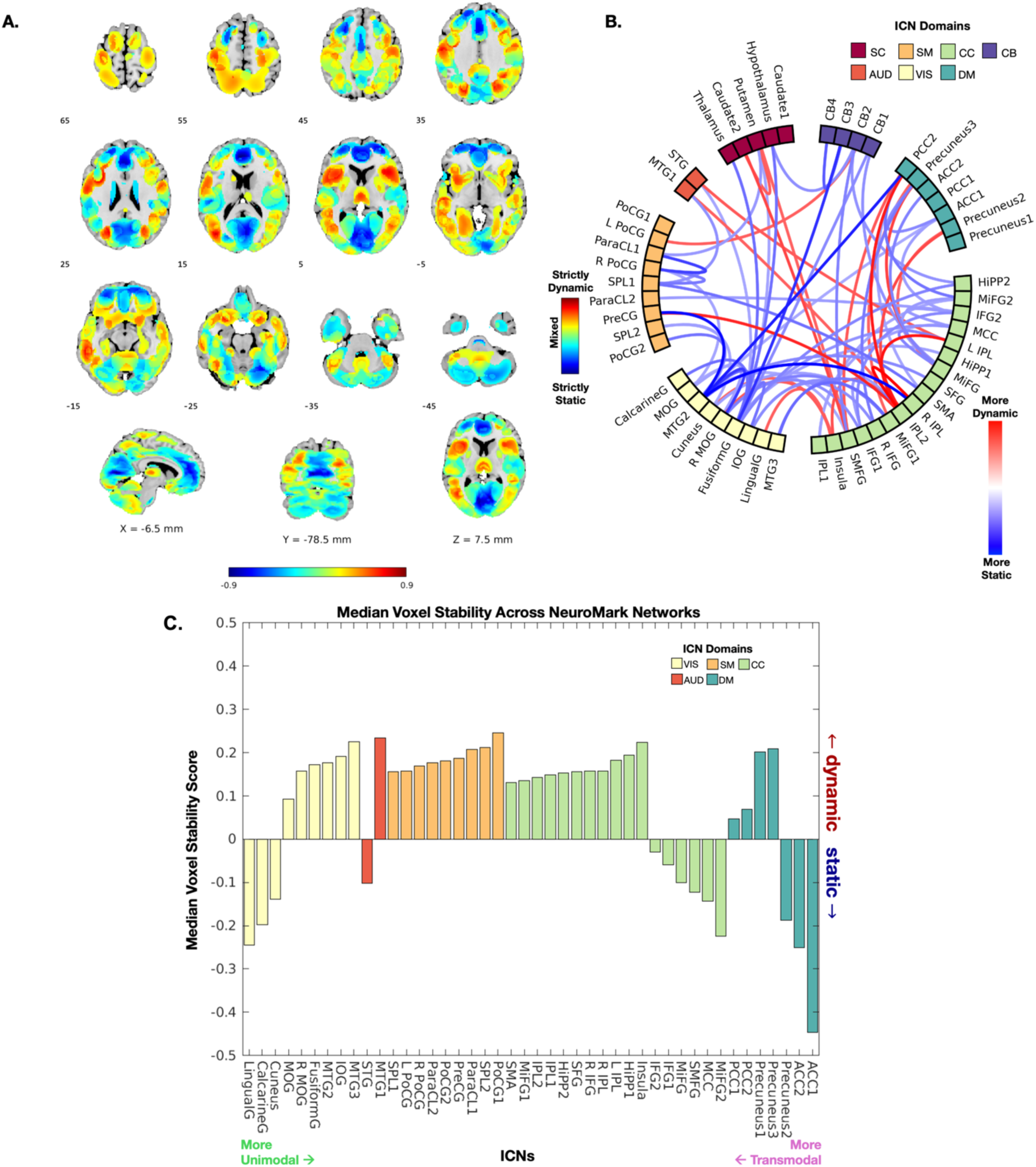
Average of Rest1 and Rest2 differential stability maps in both structural (A) and functional (B) components. Structural maps reveal organization of the brain into regions hallmarked by static (anterior cingulate, visual regions) and dynamic (sensorimotor, subcortical regions, insula) structure/function linkages. Superimposing cortical NeuroMark ICNs onto the structural map in (A) reveals alignment of static hubs in ICNs towards the extremes of unimodal/transmodal cortical hierarchy, while the majority of dynamic regions are found in intermediate regions along this gradient, as measured by median voxel value within ICNs (C).

From the structural differential stability map (Fig. 6A), we can observe the organization of the brain into clear hubs of static and dynamic structure/function linkage, as well as mixed regions that exhibit both properties. Static hubs are comprised mostly of regions in the anterior cingulate and visual regions, whereas dynamic hubs are found in the insula, sensorimotor regions, and subcortical structures. These patterns are also reflected in the differential functional component map – the insula, inferior parietal lobule, posterior cingulate, precuneus and subcortical structures including putamen and caudate are all marked by stronger edges in dynamic components, whereas visual, sensorimotor and anterior cingulate regions all serve as hubs for edges in predominantly static joint sources. We further digest the structural differential stability map by superimposing the 44 NeuroMark ICNs corresponding to cortical regions and computing the median voxel stability values (Fig. 6C). Sorting these functional domains from more unimodal to more transmodal, we observe the most highly static ICNs (stability < 0) align towards the extremes of this functional gradient, whereas the most dynamic ICNs (stability > 0), are situated in intermediary regions, including the entirety of the sensorimotor domain.

### White Matter Components Also Show Strong Links to Temporally Evolving FNC

Of the total 833 HCP subjects, 737 had processed FA maps available, thus we replicated the state-level HCP dynamic fusion experiments in this subset of subjects using white matter FA maps as the structural inputs with model order = 10. We reduced the model order relative to the GMV experiments to accommodate for the reduction in relative brain volume corresponding to the WM tracts in the FA data, and report results for GMV fusion with the same model order = 10 for ease of comparison. Results comparing the GMV and FA dynamic fusion experiments with the same model order are shown in Fig. 7. While the overarching relationship between structure and function holds in the FA data (higher stability in structural components than functional components), there are a few key differences to highlight. First, the FA experiment shows evidence for just one highly static (mean cross-fusion correlation = 0.75), with a steep drop-off to the rest as dynamic components, as opposed to a gradual decrease in cross-fusion stability and two or three “static” component maps. Second, the overall cross-fusion stability of the components is lower in the FA experiments compared to GMV. This result may suggest a stronger linkage to the changing functional manifolds in each distinct dFNC state fusion, thus leading to lowered cross-fusion stability. To assess this hypothesis, we compared the correlations between the structural and functional loading parameters for all components in the GMV and FA experiments and found that the correlations were indeed higher in the FNC + FA components (p = 0.0043, t = -2.927). These results may suggest that white matter FA components can be more susceptible to influence of changing functional inputs than GMV maps in symmetric data fusion approaches like mCCA + jICA.

**Fig. 7.**
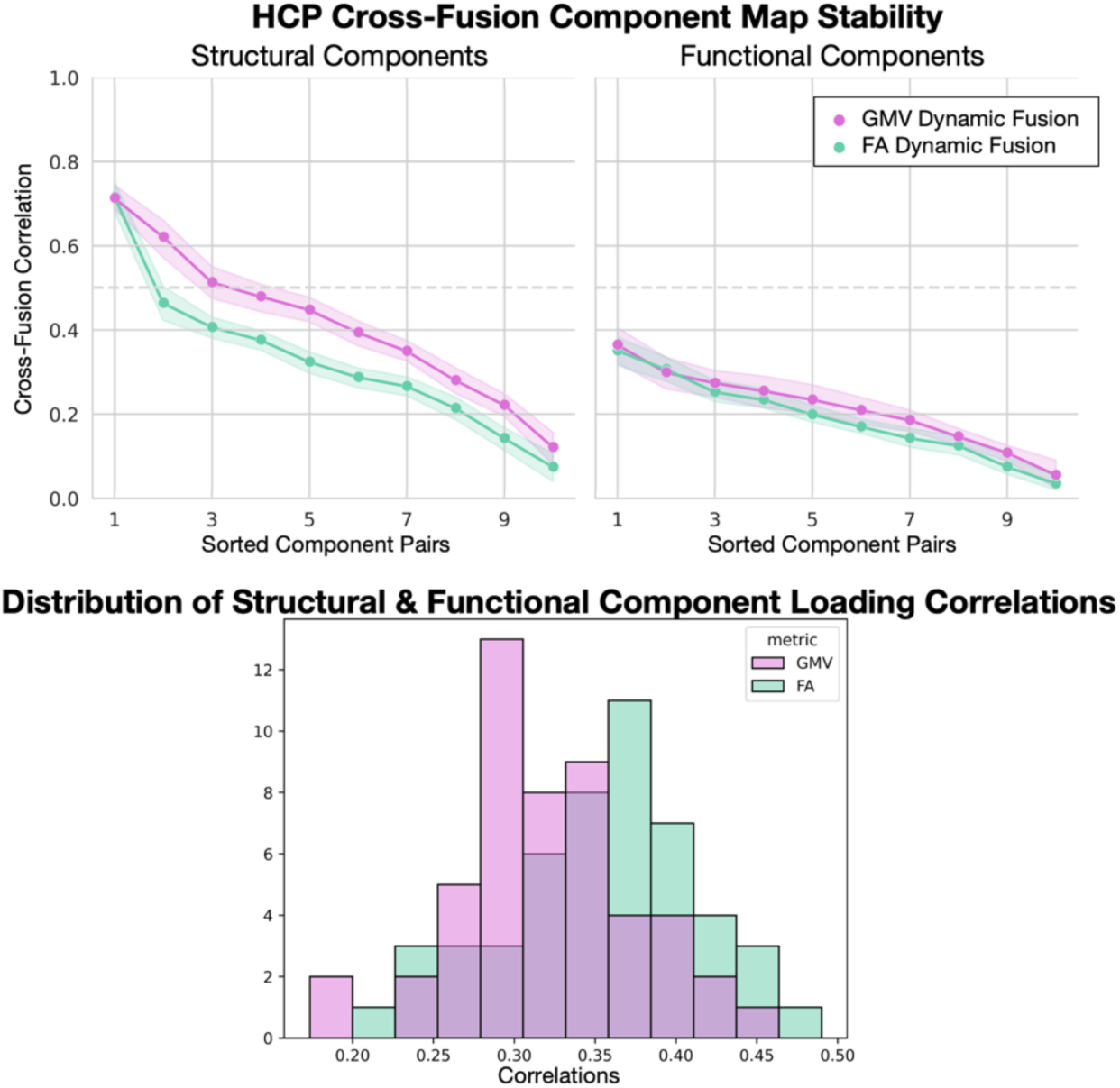
White matter (FA) components show one highly stable component, with a steep drop-off to dynamic components that show overall lower cross-fusion stability than corresponding GMV components in the same subjects (top panels). Higher correlation between structural and functional loading parameters was observed in FA fusion experiments compared to GMV fusion (bottom panel; p = 0.0043, t = -2.927).

## Discussion

Here, we propose an approach for identifying flexible, dynamic linkages between brain structure and time-varying brain function, termed dynamic fusion. Though previous studies have examined time-resolved coupling of brain structure and function (Duda et al., 2022; Iraji et al., 2019; Liu et al., 2022; Plis et al., 2018), to the best of our knowledge our work is the first to jointly estimate voxel-wise structural components as a function of time, and thus as a function of structure’s changing linkage to dynamic brain function. Our approach is fully data driven and allows both modalities (or all when 3+ modalities are used) to contribute to the fusion equally (i.e., symmetric fusion), thus enforcing fewer assumptions and enabling a broader spectrum of flexibility than recent symmetric and asymmetric fusion works in structural dynamics. The findings outlined in this paper show how multiple linked fusions can be utilized to flexibly study dynamic structure-function coupling in multimodal neuroimaging, and emphasize value of such approaches in both clinical and control populations.

We show that dynamic fusion identifies functionally adaptive neuroanatomical basis sets related to changing functional manifolds at the dFNC state-level. To fully appreciate these results, we wish to underscore that the structural inputs are identical at each independent fusion within a given experiment, thus *any change in the resultant structural component maps is driven solely by its varying linkage to the changing functional inputs* at each window/state. When applied in resting state data, we find only a few GMV components are consistently derived with high correspondence when jointly estimated with changing functional manifolds (Fig. 2), while the vast majority show evidence of more significant functional influences. These results held across numerous experiments encompassing different data sets (HCP vs. FBIRN), subject cohorts (CON vs. SZ), dynamic FNC approaches (SWPC vs. FBC), and even structural inputs (GMV vs. FA maps). Taken together, these results exemplify mCCA + jICA’s capacity to identify truly joint linkages in the data to produce structural components that are indeed functionally influenced. Moreover, our findings underscore the importance of enabling flexible linkages between structure and function, especially as a function of the dynamic brain changes over time. Without the natural flexibility afforded by the mCCA + jICA algorithm, as well as the flexibility imparted by conducting multiple linked fusions as the novelty of the approach taken in this work, the nuanced joint structure-function relationships would have remained hidden in the data (Sui et al., 2013).

In addition to the cross-fusion stability of components related to changing cognitive demands, we found that a large portion (40/73) of dynamic components showed significant group differences between SZ and controls in the FBIRN resting state experiments, with the most significant group difference belonging to a highly dynamic component (Fig. 5). Based on this result, we can conclude functionally influenced structural components can indeed capture the changes in the relationship between structure and dynamic brain function of clinical and diagnostic relevance. By assessing the full structural/functional loading parameter space we also conclude that dynamic components show stronger separability along both structural and functional loading parameter axes than comparable static components, suggesting dynamic components do, indeed, represent a more harmonious joint influence of structure and function on the joint source overall and thus better capture the complex interplay of structure and function underpinning both clinical and control populations.

Moreover, by applying dynamic fusion in a set of healthy control data from the HCP cohort we generated a map of global differential stability, organizing the brain into regions of static and dynamic functional linkages. We found many static hubs aligned to the extremes of the unimodal (i.e., the cuneus, calcarine gyrus and lingual gyrus within the visual network) – transmodal (i.e., anterior cingulate cortex within the default mode network) hierarchy of cortical structures, whereas dynamic hubs were comprised of intermediary regions along this gradient (i.e., insular cortex, frontoparietal/sensorimotor regions) (Margulies et al., 2016). The significance of this result is two-fold: 1) it upholds and closely mirrors results of a recently published study which utilized multilinear regression to evaluate time-resolved structure-function coupling in atlas-derived brain networks (Liu et al., 2022), and 2) further underscores the neurobiological relevance of sources derived via our dynamic fusion approach.

It is thought that the unimodal-transmodal gradient reflects a similar gradient in connection lengths across the cortex (Oligschläger et al., 2019; Sepulcre et al., 2010). Thus, it is possible that the stability observed at the poles of this gradient are also a reflection of connection lengths – speculatively, this could stem from the ease of maintaining shorter connections and the importance of stabilizing critical long-range connections. In their work, (Liu et al., 2022) found this same U-shaped relationship between stability and connection length, with highest stability at the shortest and longest extremes and increased dynamicity for those networks with mid-range connection length. Our results largely align with this finding, and while not a 1:1 mapping of dynamicity to this cortical hierarchy, they do warrant future exploration into connection lengths as a facet of structure-function dynamics.

One of the clearest dynamically linked structure-function hubs in both our study and (Liu et al., 2022) is the insula. The insula is a key region in the ventral attention/salience network that dynamically fields and coordinates communication of both unimodal (e.g., sensory) and transmodal (e.g., default mode) regions (Ghaziri et al., 2017; Tian & Zalesky, 2018). Taken together with our results, we echo the speculations presented in (Liu et al., 2022) and suggest that dynamic structure-function coupling exhibited within the insular cortex could reflect flexible integration and transmission of various neural signals though this complex cortical hub. Thus, states that result in dynamic structural components with peaks in insula could be thought of as “salience” states in which attention is being directed or integrated within the cortex.

This work represents a reduced-case feasibility study of a wider dynamic fusion framework (Fig 1). Our results clearly demonstrate the utility of estimating flexible linkages between structure and dynamic brain function, and thus motivate the extension of dynamic fusion to a more advanced model. In our next step, we plan to extend this work into a unified fusion framework that will jointly estimate multiple structural basis sets at once based on unique linkages between structural and temporally evolving functional inputs. In addition to the flexibility of coupling between neuroanatomical measures and dynamic connections between functional networks, we may also consider incorporating flexibility in the structural definitions of the functional networks themselves. Future work could employ dynamic spatially constrained ICA (Duda et al., 2023; Iraji et al., 2019) to extract individualized time-varying functional entities with flexible spatial maps at each window in the dynamic analysis. Such an analysis could be done using the original 53 ICNs defines in the Neuromark_1.0 template, or be expanded to the multiscale Neuromark template (http://trendscenter.org/data), which defines 105 ICNs derived across multiple spatial and temporal scales (Iraji et al., 2023), further increasing the precision with which we can investigate dynamic structural and functional coupling in a data-driven manner.

Other future directions may involve extending the analyses described here by applying 3-way dynamic fusion to enable deeper investigation into the connection of both gray matter and white matter structure to temporally evolving brain function, as well as how hidden linkages between the structural modalities affect the joint estimation of the structural components. Interestingly, one result in our study suggested that white matter FA components were potentially more susceptible to functional influence in the joint fusion than GMV components in the same subjects (Fig. 7). Future work will be needed to replicate these results and further investigate the nature of white matter structure/gray matter function linkages. A newly developed ICA-based approach has enabled joint estimation of linked structural and functional connectivity from dMRI and fMRI data (Wu et al., 2015; Wu & Calhoun, 2023). Future work could leverage this approach under the dynamic fusion framework further address time varying structure-function linkages between white matter and FNC. Thus far, the analysis of brain function is largely focused on grey matter regions, leaving functional activity and connectivity within white matter regions relatively understudied. In fact, in data-driven analyses, functional components whose structural maps reside largely within white matter are often discarded as noise components. Exploration of the role of these white matter functional components in the context of whole brain function and cognition is currently underway. Future work including white matter functional components would further contextualize the investigation into dynamic structure-function coupling with tissue-level specificity on both ends.

The results presented in this work are subject to various limitations. First, the functional inputs into our fusion experiments were all derived from a single functional template, NeuroMark 1.0. This template is well established and validated for capturing both subject-level variations and strong group-level correspondence (Du et al., 2020), however the networks can be considered relatively coarse-grained owing to the relatively small template size of 53 ICNs. Recently, more fine-grained templates have been developed (Iraji et al., 2023) and refined (Jensen et al., 2024) that define functional ICNs along multiple spatial scales, as well as several templates developed for specific age ranges across the lifespan (Fu et al., 2024). Future work may consider a comparative analysis between dynamic fusions utilizing different spatial scale templates. Similarly, the specifications of the subject cohorts used within our analyses should also be considered. The history of antipsychotic and other medications was not explicitly tested within our experiments in the FBIRN SZ cohort, thus interpretation of our results should be considered in that context. Beyond this, SZ is known to show significant overlaps, both functionally and structurally, with other conditions including bipolar disorder and schizo-affective disorders (Clementz et al., 2022), so specificity of our findings to SZ in the context of other conditions necessitates further study. In the HCP experiments, the study cohort represents a relatively narrow age range (22 – 37 years), thus results may not generalize outside of young adulthood. Replication in cohorts with a wider age range is warranted. Moreover, by applying dynamic fusion in additional datasets and subject cohorts, we can more readily understand the generalizability of the results reported here, as well as explicitly test the relationship between static/dynamic components and connection lengths across the unimodal/transmodal principal functional gradient.

In conclusion, we present an approach for dynamic fusion of multimodal neuroimaging data as a method for studying time-varying structure-function coupling. Components derived from our dynamic fusion approach exhibit strong cognitive, clinical and neurobiological correlates, underscoring the importance of retaining rich temporal dynamic information in studies of structure-function linkage.

Overall, our results support the claim that patterns of structure-function coupling are both regionally heterogenous and temporally varying and provide an important next step in developing a model of dynamical linkage between brain function and structure.

## Data and Code Availability

Details for the availability of the FBIRN dataset used in this paper can be found here: https://www.nitrc.org/projects/fbirn/.

Details for the availability of the HCP dataset used in this paper can be found here: https://www.humanconnectome.org/study/hcp-young-adult.

Code and documentation for running mCCA + jICA fusion can be found in the FIT toolbox here: https://trendscenter.org/software/.

## Author Contributions

This study was conceived and designed by MD & VDC. Coding, analysis, visualization of results, and initial drafting of manuscript was performed by MD. All authors participated in interpretation of results and editing the final manuscript.

